# Membrane cholesterol regulates TRPV4 function, cytoskeletal expression, and the cellular response to tension

**DOI:** 10.1101/2020.12.01.406850

**Authors:** Monika Lakk, Grace F. Hoffmann, Aruna Gorusupudi, Eric Enyong, Amy Lin, Paul S. Bernstein, Trine Toft-Bertelsen, Nanna MacAulay, Michael H. Elliott, David Križaj

## Abstract

Despite the association of cholesterol with debilitating pressure-related diseases, its role in mechanotransduction is not well understood. We investigated the relationship between mechanical strain, free membrane cholesterol, actin cytoskeleton, and activation of stretch-activated TRPV4 (transient receptor potential vanilloid isoform 4) channel in human trabecular meshwork (TM) cells. Physiological levels of cyclic stretch resulted in time-dependent decreases in membrane cholesterol/phosphatidylcholine ratio and upregulation of stress fibers. Depletion of free membrane cholesterol with m-β-cyclodextrin (MβCD) augmented TRPV4 activation by the agonist GSK1016790A, swelling and strain, with the effects reversed by cholesterol supplementation. MβCD increased membrane expression of TRPV4, caveolin-1 and flotillin. Caveolin-1 antibody partially precipitated a truncated ∼75 kDa variant whereas the majority of TRPV4 did not colocalize or interact with caveolae or lipid rafts, indicating that TRPV4 is mainly localized outside of cholesterol-enriched membrane domains. MβCD induced currents in TRPV4-expressing *Xenopus laevis* oocytes. Thus, while the membrane C/P ratio reflects the biomechanical milieu, trabecular transduction of mechanical information is modulated by the membrane cholesterol content. Diet, cholesterol metabolism and mechanical stress might modulate the conventional outflow pathway and intraocular pressure in glaucoma and diabetes.

## Introduction

Conversion of sensory information into electrical and chemical signals in eukaryotic cells is modulated by unesterified cholesterol, a planar 27-carbon polycyclic molecule that constitutes ∼20% of the total mass of membrane lipids (1, 2). Its intercalation into phospholipids reduces the motion of hydrocarbon chains, affects surface charge, and promotes membrane stiffness while keeping the membrane fluid (3–5). Cholesterol is also a precursor for steroid hormones and regulates stereospecific interactions with transmembrane channels, transporters, and enzymes within membrane-bound glycosphingolipid-rich protein complexes (lipid rafts; 6) that serve as organizing hubs for intracellular signaling pathways, membrane trafficking, cytoskeleton, and cell-extracellular matrix (ECM) interactions (1, 2, 7, 8). Alterations in free membrane cholesterol and lipid raft density provoke positive and negative effects on membrane channels, with little consensus about the molecular mechanisms (9–11).

Vertebrates maintain cholesterol levels within a narrow range, with deficits or oversupply often deleterious for health (5, 6). Defects in cholesterol regulation contribute to atherosclerosis, myocardial injury, diabetic and vascular dysfunctions (12–14) and increase the risk for progression of neurodegenerative diseases such as primary open-angle glaucoma (POAG), a prevalent blinding disease (15, 16). Patients with high levels of serum cholesterol have increased likelihood for elevated intraocular pressure (IOP), with glaucoma linked to multiple genes involved in cholesterol metabolism (17, 18). IOP is dynamically regulated by the trabecular meshwork (TM), a multilayered tissue composed of smooth-muscle-like cells with mechanosensitive and contractile functions that control the outflow of aqueous humor from the anterior eye (16, 19–21). In response to mechanical stress and glaucoma, TM cells upregulate cytoskeleton and secretion of ECM, with attendant increases in cell contractility and stiffness producing a suppression of aqueous outflow and elevation in IOP (22, 23, 36, 37). It is not known, however, how TM membrane lipid composition is affected by the biomechanical milieu, and whether changes in membrane stiffness impelled by free membrane cholesterol affect TM mechanosensing, intracellular signaling and cytoskeletal organization.

We recently identified the mechanosensitive Transient Receptor Potential Vanilloid 4 (TRPV4) channel as a principal regulator of mechanically induced signaling in mouse and human TM (20, 24, 25). This ubiquitous nonselective cation channel transduces membrane strain, shear flow, swelling, temperature and polyunsaturated fatty acids into calcium signals (21, 24, 26, 27). Its sequence appears to have co-evolved with enzymes linked to cholesterol biosynthesis pathways (28) and studies in endothelial cells suggested that TRPV4 may be trafficked to cholesterol- and caveolin1 (Cav-1)-enriched rafts (29) to modulate vascular flow (30). This study’s objective was to determine the mechanistic link between mechanotransduction, TRPV4 signaling and cholesterol in human TM cells and in an oocyte overexpression system. We found that the lipid composition of the TM membrane reflects the cells’ history of exposure to mechanical stress. Membrane TRPV4 was mainly located in non-raft regions, as indicated by the lack of interaction with Cav-1 and the caveolar accessory protein flotillin. Manipulation of free membrane cholesterol regulated TRPV4 activation, and thereby its role in calcium homeostasis and cytoarchitecture. Hence, the TM capacity to homeostatically regulate the aqueous outflow pathway might be under dynamic, compounded influence of diet, the biomechanical milieu and systemic/local cholesterol.

## Results

### Tensile stretch regulates the lipid content of the TM membrane

The mechanical properties of the lipid bilayer can change under tension (31), but it is not known whether the membrane lipid content is shaped by the biomechanical milieu. To test this, we determined the changes in cholesterol content and the C/PC (cholesterol/phosphatidylcholine (PC)) molar ratio following the stimulation of cells with cyclic mechanical strain. Primary human TM cells were isolated from healthy donors (24; 25), plated on ECM (Collagen I)-coated membranes and stimulated for 1 or 3 hours with radial stretch (0.5 Hz; 6% elongation). Membrane cholesterol and phosphatidylcholine levels were measured with GS/MS and a fluorometric PC assay. Normalized relative to unstimulated controls, the stretched samples showed time-dependent decrease in membrane cholesterol content to 0.58 ± 0.089 (1 hour; N = 3; P < 0.01) and 0.30 ± 0.02 (3 hours; N = 3; P < 0.001) (Fig. 1A & B) whereas the PC content increased to 1.32 ± 0.02 (1 hour; N = 3; P < 0.0001) and 1.58 ± 0.03 (3 hours; N = 3; P < 0.01), respectively (Fig. 1C). The membrane C/PC ratio (control, C/PC = 1) thus decreased to 0.44 following 1-hour, and 0.19 after 3 hours of mechanical stimulation. Its sensitivity to the mechanical strain suggests that mechanical properties of the cell membrane reflect the history of the exposure to the biomechanical environment.

**Figure 1.**
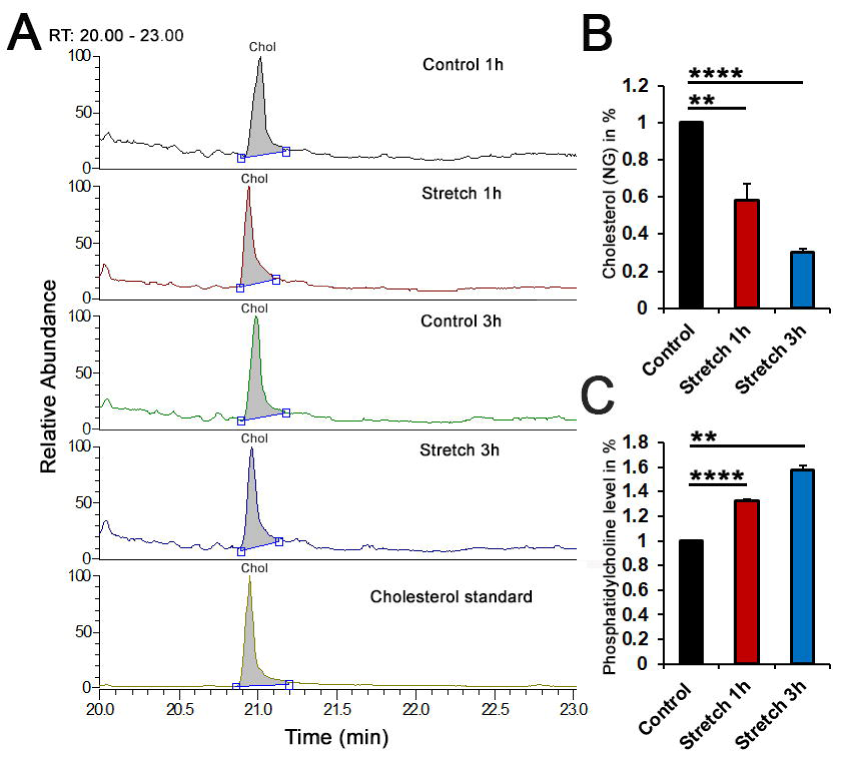
The C/P ratio in TM membranes is stretch-dependent. (A) Representative chromatograms and (B) Normalized and averaged cholesterol GS/MS data in control, 1 h and 3 hour stretched primary TM cells, (C) Normalized and averaged phosphatidylcholine (PC) fluorometric data in control, 1 h and 3 hour stretched primary TM cells. Stretch induced a time-dependent decrease in membrane cholesterol level, which was concomitant with increasing content of membrane P.C. N = 3; **, P < 0.01; ****, P < 0.0001.

### Reduction in free membrane cholesterol is associated with differential upregulation of ion channels

To determine whether stretch-dependent modulation of membrane cholesterol affects mechanosensing and transduction, cells were exposed to methyl-beta-cyclodextrin (MβCD), a water-soluble cyclic oligosaccharide that encapsulates hydrophobic membrane cholesterol residues and has been widely used to characterize cholesterol-dependence of ion channels (32). Filipin, a fluorescent polyene antibiotic (33) was used to visulize endogenous unesterified cholesterol within lipid rafts (34). 60 min incubation with MβCD (10 mM) reduced filipin-β positive puncta by 75.63% ± 2.77% (P < 0.0001) (Fig. 2A,B and D), with the cells remaining viable and responsive to physiological stimuli over the duration of a typical experiment (∼1-3 hours). Cholesterol:MβCD supplementation (1:10) (35) increased the number of filipin-positive puncta by 6.17 ± 0.67-fold (Fig. 2D), suggesting that cholesterol facilitates the formation of raft domains.

**Figure 2.**
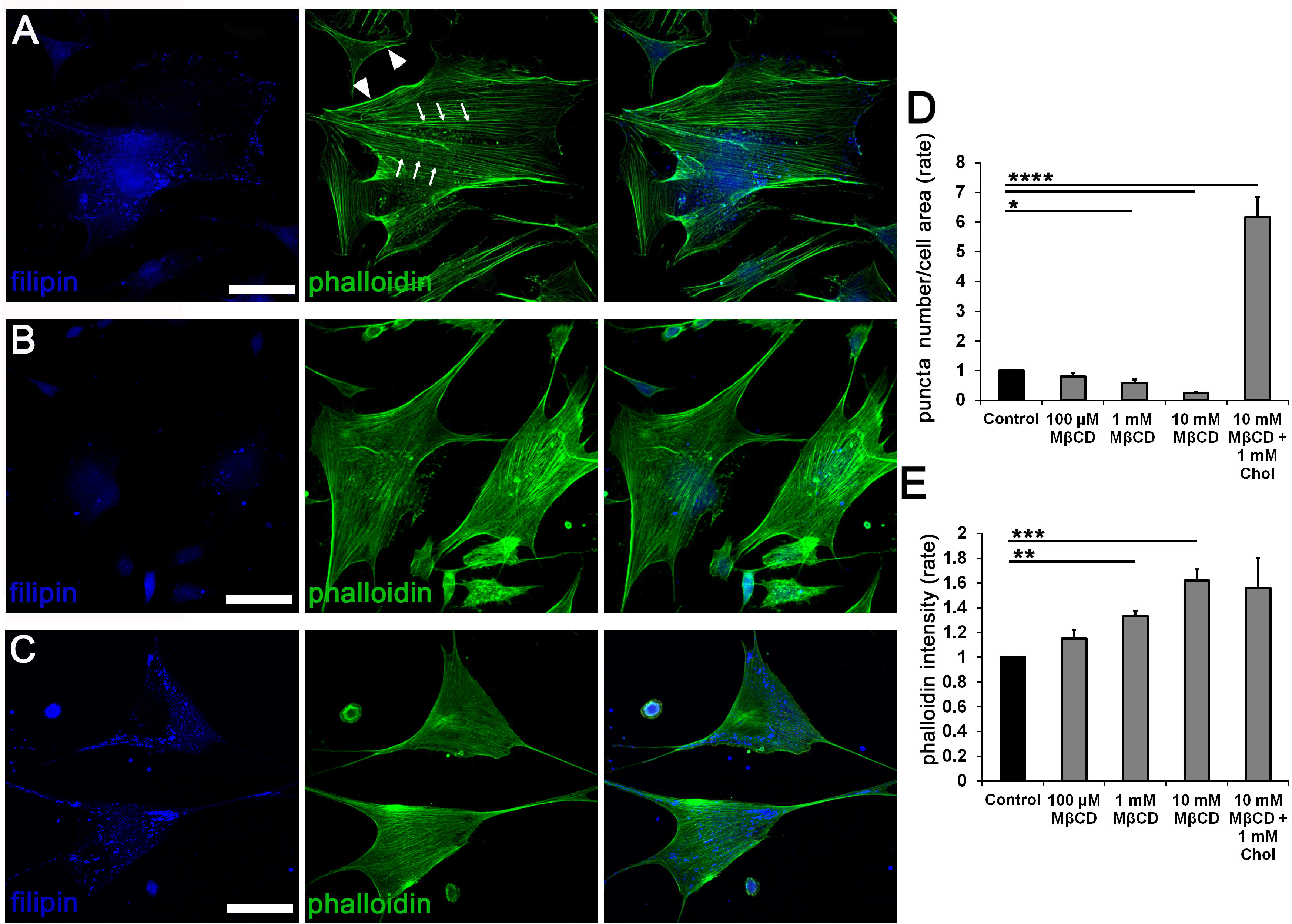
Cholesterol depletion promotes formation of actin stress fibers. Double-labeling for F-actin (phalloidin-Alexa 488 nm) and filipin (405 nm). (A) Untreated preparations show typical stress fiber organization dotted by lipid rafts. (B) 1-hour incubation with MβCD results in dissolution of filipin^+^ puncta and upregulation of phalloidin-actin fluorescence. (C) Saturated 1:10 admixture of cholesterol (1 mM) and MβCD (10 mM) increased the number of filipin puncta. (D-E) Averaged data for experiments shown in A-C (N = 3-4). Scale bar: 50 µm. *, P < 0.05, **; P < 0.01, ***, p < 0.001 and ****, p < 0.0001.

### Reduction in free membrane cholesterol is associated with upregulated expression of F-actin

Mechanical stability of cells is maintained through interactions between the cell membrane, the cytoskeleton and the ECM. Cortical actin (arrowheads in Fig. 2A) and ventral stress fibers (arrows) provide structural integrity and mediate cell – ECM contacts and contractility in unstimulated TM cells but may be reinforced in response to mechanical stress (19, 25, 36). We tested whether altering membrane stiffness through cholesterol depletion/enrichment impacts cytoskeletal architecture in cells labeled with phalloidin-actin Alexa 488. 1-hour exposure to 0.1 – 10 mM MβCD was associated with dose-dependent increases in stress fiber fluorescence. 10 mM cyclodextrin produced 55.8 + 9.5% increase in the F-actin signal (Fig. 2B & E) (N = 4; P < 0.001) (N = 4; P < 0.001) whereas supplementation with saturated 1:10 (1 mM) mixture of cholesterol and MβCD did not affect actin upregulation (Fig. 2C & E) despite the sizeable increase in the number of filipin^+^ puncta (Fig. 2C &D). Thus, actomyosin organization in TM cells does not appear to require lipid rafts.

### Cholesterol depletion facilitates agonist-induced TRPV4 activation

TRPV4, a polymodal calcium-permeable channel that recently emerged as potential regulator of TM pressure, strain and volume sensing and conventional outflow (20, 24, 25) contains putative cholesterol recognition motifs within Loop4-TM5 (28). Its sensitivity to cholesterol modulation was investigated in cells loaded with the Ca^2+^ indicator dye Fura-2-AM and stimulated with the agonist GSK1016780A (GSK101) before and after exposure to MβCD. As previously shown (24, 25), GSK101 (25 nM) induced reversible increases in [Ca^2+^]_i_ (Fig. 3A). MβCD increased peak amplitude of GSK101-evoked [Ca^2+^]_i_ elevations from 0.5 ± 0.04 (n = 39; N = 6) to 1.06 ± 0.13 (n = 53; N = 6; P < 0.0001) (∼100% increase) without affecting baseline [Ca^2+^]_i_. Agonist stimulation under cholesterol-enriched conditions produced a small but significant increase in the average GSK101 response (0.74 ± 0.05; n = 44; N = 5; P < 0.001).

**Figure 3.**
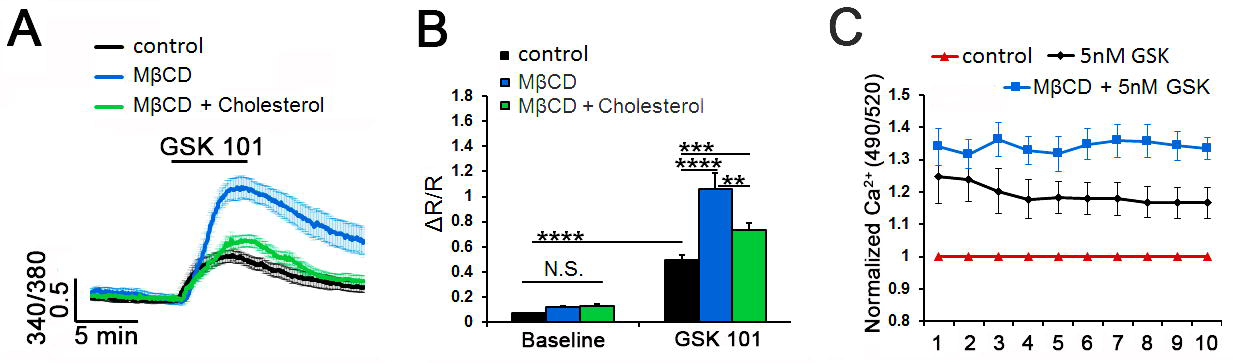
Cholesterol depletion increases the amplitude of TRPV4 agonist-induced Ca^2+^ signals. (A-B) Ratiometric signals in Fura-2 AM-loaded cells. (A) GSK101-induced elevations are increased in Mβ CD (blue bar) augmented, whereas MβCD:cholesterol (green bar) reduced the amplitude of agonist-induced fluorescence. (C) Fluorimetry, cell populations in 96 wells. 5 nM GSK101 increased Fluo-4 fluorescence. Its effect was facilitated (∼ 12%) by βCD (N = 4). **, P < 0.01; ***, P < 0.001; ****, P < 0.0001; N.S., non-significant.

Spectrophotometry was used to assess the effects of cholesterol depletion on the TM population response. 5 nM GSK101 produced a 24.8% ± 8.31% increase in the Fluo-4 fluorescence signal (N = 4 independent experiments; P < 0.05; Supplementary Fig. 1A). MβCD potentiated the response by ∼ 12%, resulting in 36.27% ± 5.18% increase compared to the baseline (P < 0.05) (Fig. 3C). A similar facilitatory effect was observed with 25 nM GSK101 (Supplementary Fig. 1A). These data suggest that cholesterol suppresses agonist-induced TRPV4 activation.

### Cholesterol modulates swelling -induced calcium signaling

Originally identified as a regulator of cellular swelling (38, 39), TRPV4 functions as a real-time readout of cell volume changes (40) with osmoregulatory functions in neurons, glia, epithelial and endothelial cells (41–46). To determine the effect of raft disruption on the TM swelling response, we exposed the cells to hypotonic stimuli (HTS) in the presence/absence of MβCD. Consistent with TRPV4 activation, TM cells responded to 140 mOsm HTS with significant (0.44 ± 0.06 Y; n = 24; N = 4; P < 0.0001) increases in [Ca^2+^]_i_ (Fig. 4A & B). Depletion of cholesterol approximately doubled the peak response amplitude to 0.87 ± 0.08 (n = 27; N = 4; P < 0.001), with the facilitatory effect inhibited by cholesterol supplementation (0.43 ± 0.04; n = 30; N = 4; P < 0.001) (Fig. 4A & B). Spectrophotometry similarly showed dose-dependent [Ca^2+^]_i_ increases in response to HTS (in case of 55% HTS: 1.14 ± 0.03; N = 3), and augmentation by cholesterol removal (to 1.21 ± 0.022; N = 3) (Fig. 4C & Supplementary Fig. 1B) (P < 0.05).

**Figure 4.**
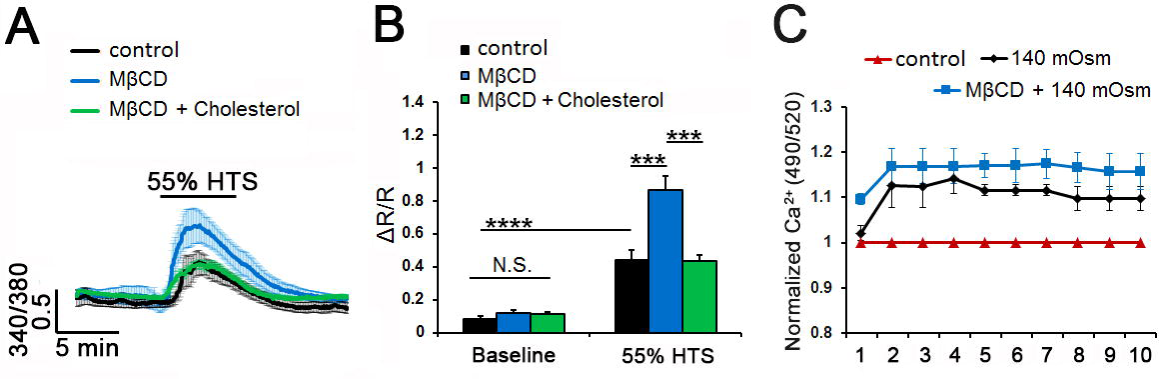
Cholesterol depletion facilitates HTS-induced Ca^2+^ signals. (A-B) Cytosolic Ca^2+^ responses in Fura-2 AM-loaded cells. (A) HTS induced [Ca^2+^]_i_ elevations in representative control, Mβ Mβ and Mβ D:cholesterol -treated samples (n = 8-10) (B) Averaged data from A. CD:cholesterol (green bar) reduced HTS-induced [Ca^2+^]_i_ increases. (C) Fluorimetry. 55% HTS (140 mOsm) increased Fluo-4 signals. This effect was facilitated by MβCD (N = 3). ***, P < 0.001; ****, P < 0.0001; N.S., non-significant.

### Cholesterol modulates the TM response to membrane strain

We next investigated whether cholesterol depletion impacts the transduction of cyclic mechanical stretch, which mirrored strains impelled on ECM beams by IOP fluctuations (47–49) that are transduced partly via TRPV4 (20, 25). Cells were stimulated with periodic, calibrated displacements/relaxations of the collagen IV–coated substrate (1 Hz; 10% elongation, 15 minutes). Cyclic stretch reversibly elevated cytosolic [Ca^2+^]_i_ to 0.21 ± 0.02 (n = 21; N = 6; P < 0.0001). The effect was augmented by cholesterol depletion to 0.71 ± 0.04 (n = 77; N = 5; P < 0.0001) and reduced by cholesterol supplementation to 0.38 ± 0.05 (n = 37; N = 6; P < 0.0001) (Fig. 5A & B). Simvastatin (10 μM), an inhibitor of 3- hydroxy-3-methylglutaryl (HMG) coenzyme A reductase, the rate-limiting enzyme responsible for endogenous production of cholesterol, induced a small, non-significant decrease in the number of fillipin^+^ puncta (Supplementary Figs. 2A & B). Accordingly, exposure to the statin did not affect the amplitude or kinetics of stress-induced [Ca^2+^]_i_ elevations (0.16 ± 0.01; n = 12; N = 2) (Supplementary Figs. 2C & D).

**Figure 5.**
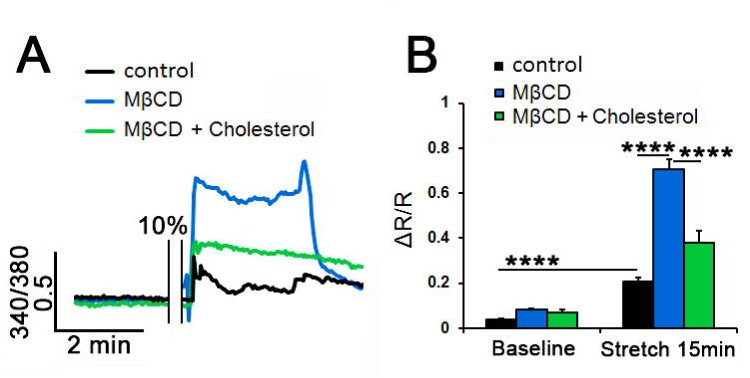
Membrane cholesterol modulates the cyclic stretch induced intracellular calcium responses in TM cells. (A) Representative traces and (B) Averaged results for 15 min cyclic stretch of control (black trace, bars), MβCD-treated (10 mM, blue trace, bar) and MβCD:cholesterol-treated (green trace, bar) TM cells. Stretch-evoked [Ca^2+^]i responses are modulate by membrane cholesterol (n = 21-77; N = 5-6). ****, P < 0.0001.

Changes in membrane cholesterol content have been shown to influence cell surface tension and actomyosin assembly (89). To test the effect of membrane cholesterol on stress fibers, F-actin was labeled with fluorescent phalloidin following exposure to cyclic stretch in the presence/absence of MβCD and MβCD:cholesterol. As shown previously, stretch alone augmented phalloidin-actin fluorescence (to 1.32 ± 0.05; N = 3; P < 0.001) (Fig. 6A & C). Interestingly, this was associated with a significant decreased in the density of filipin puncta (to 0.76 ± 0.05; N = 3; P < 0.01) (Fig. 6A panel ii & B), suggesting that mechanical stress indluences formation of lipid rafts. As shown in Figure 2, MβCD increased the F-actin signal (1.56 ± 0.09; N = 3; P < 0.0001) while reducing filipin fluorescence to ± 0.03 (N = 3; P < 0.0001). MβCD treatment of stretch-exposed cells did not further affect filipin fluorescence (0.26 ± 0.02; N = 3) (Fig. 6A & B) whereas stress fiber fluorescence showed a significant 42.6% ± 21.6% increase over the effect of stretch alone (N = 3; P < 0.05) (Fig. 6C). These data suggest that stretch and cholesterol depletion have additive effects on facilitating TM F-actin expression, and that stretch functions as a negative regulator of raft assembly.

**Figure 6.**
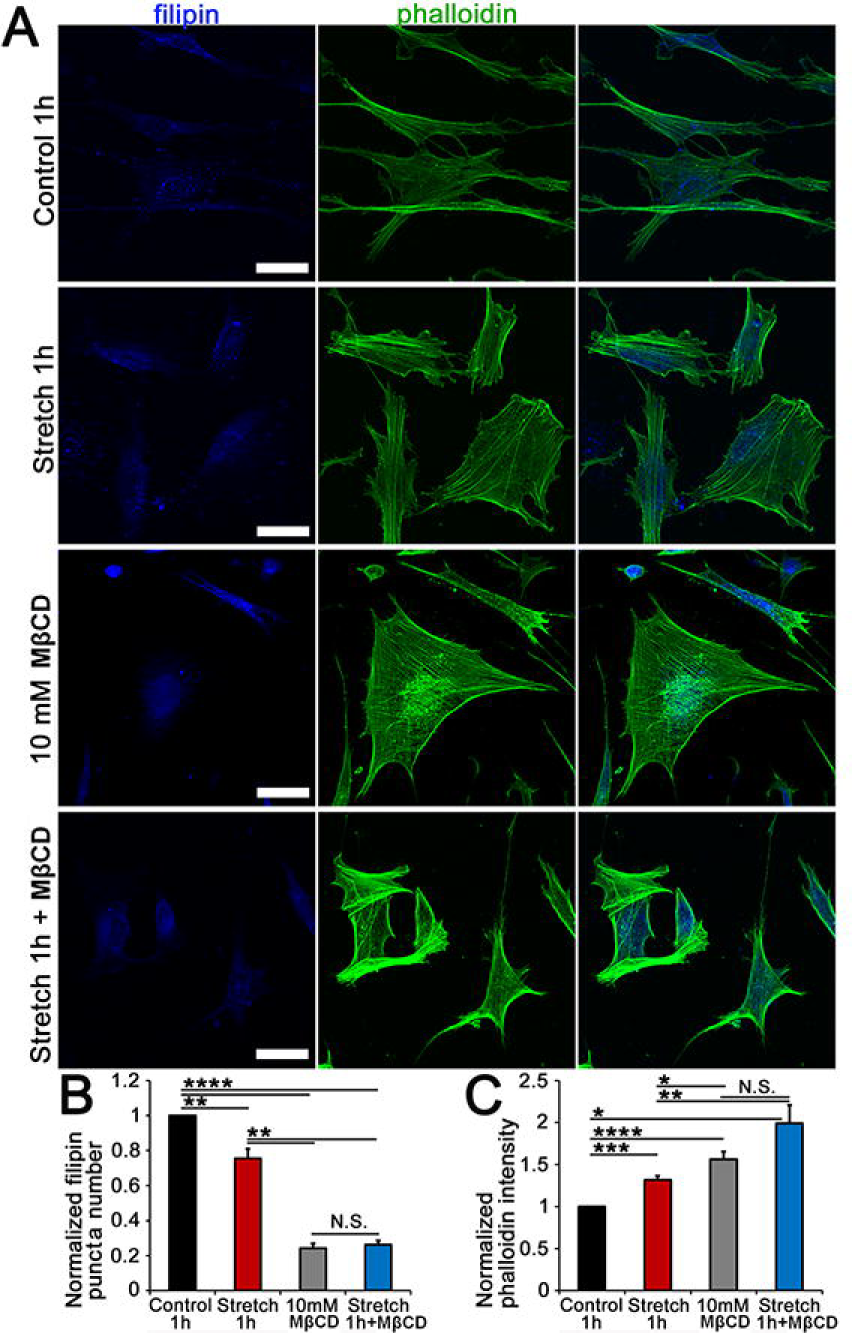
Membrane cholesterol content regulates stretch-dependence of cytoskeletal remodeling. (A) TM cells were stretched in the presence/absence of MßCD and MßCD:cholesterol, and labeled for filipin (405 nm) and phalloidin-Alexa 488 nm in control, 6% stretch (0.5 Hz, 6%, 1 hour), MßCD and MßCD + stretch samples. (B) Averaged and normalized puncta number. Filipin is significantly reduced by stretch and MßCD alone. (C) Stretch and MßCD significantly facilitate F-actin fluorescence. Combined stimulation results in an additional ∼30% increase in stress fiber signal. (N = 3-4) *, P < 0.05; **, P < 0.01; ***, P < 0.001; ****, P < 0.0001.

### Membrane cholesterol regulates TRPV4 expression

Depending on the cell type and protein isoform, cholesterol-modulating agents potentiate or inhibit TRP channel trafficking (50–52). To determine the effect of membrane cholesterol manipulations on channel expression, cells were fixed and labeled with a validated anti-TRPV4 antibody (42; 101). 1 hour incubation with Mβ produced a 4.49 ± 1.07-fold increase in the number of TRPV4-ir puncta (N = 3; P < 0.05) (Fig. 7B & D). Cholesterol enrichment abrogated this effect (0.73 ± 0.28; N = 3; P < 0.05) (Fig. 7C & D).

**Figure 7.**
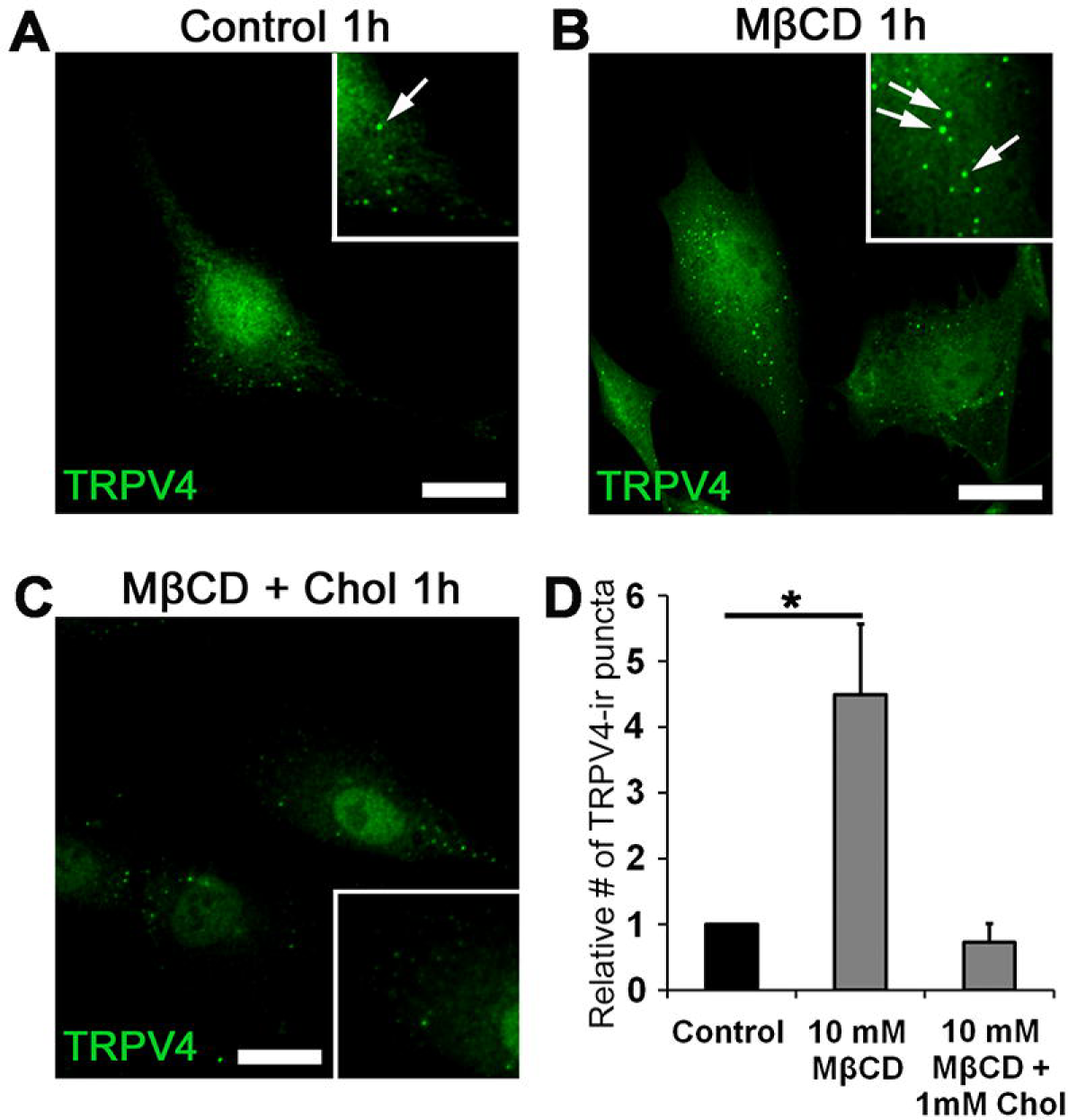
Cholesterol depletion increases the number of TRPV4-ir puncta. TRPV4 immunolabeling of primary TM cells. Representative examples of (A) Control, (B) 1-hour treatment with MßCD and (C) 1-hour treatment with MβCD:cholesterol. Inset: zoomed-in regio with TRPV4-ir puncta (arrow); (D) Summary of 3 independent experiments, normalized for control cells. The number of TRPV4 puncta is upregulated after incubation with MßCD. *, P < 0.05.

### TRPV4 current in heterologously expressing oocytes is potentiated by MβCD

The opening of the TRPV4 channel pore in mammalian cells reflects a multitude of simultaneous inputs that include mechanical stressors, temperature, polyunsaturated fatty acids and accessory binding proteins (45, 53, 54). To isolate the effects of cholesterol on the channel from auxiliary proteins and intracellular signaling within TM cells, we determined the cholesterol-dependence of TRPV4 currents in the *Xenopus laevis* expression system that has been previously used to study the effects of cholesterol depletion (55) and TRPV4 signaling (40, 42, 46). Because oocyte viability is compromised at mM MβCD concentrations (55), the experiments were conducted using 50 μM MβCD.

Membrane currents in uninjected control oocytes and TRPV4-expressing *Xenopus* oocytes were monitored by conventional two-electrode voltage clamp. A voltage step protocol demonstrated small tonic currents in TRPV4-expressing oocytes (Fig. 8A, upper left panel) compared to those obtained in uninjected oocytes (Fig. 8A, lower left panel), summarized in Fig. 8B and inset, N = 9. Transmembrane currents of uninjected oocytes were undisturbed by MβCD exposure, whereas those of the TRPV4-expressing oocytes were enhanced approximately 10-fold (from 156 ± 42 nA to 1466 ± 475 nA, N = 9, P = 0.014) by cholesterol depletion (Fig. 8A-B). These data indicate that membrane cholesterol suppresses tonic TRPV4 activity, with depletion thereof enhancing the TRPV4-mediated membrane currents.

**Figure 8.**
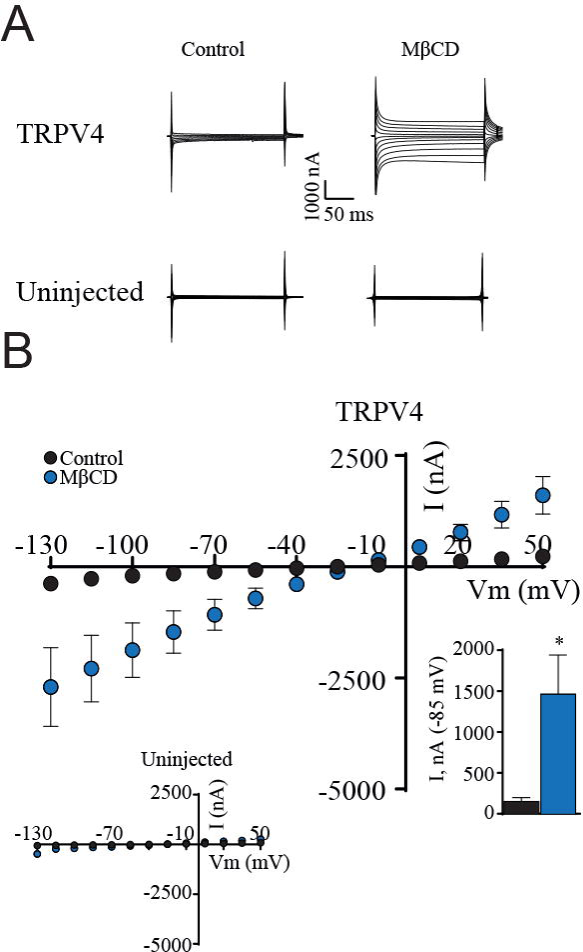
Cholesterol depletion enhances TRPV4-mediated membrane currents in *Xenopus* oocytes. (A) Representative current traces from TRPV4-expressing oocytes and uninjected control oocytes in control solution or after 45 min exposure to 50 μM MβCD. (B) I/V β curves of TRPV4-expressing oocytes exposed to control solution or MβCD, with uninjected oocytes in inset. Summarized currents obtained at -85 mV demonstrated in lower inset. The magnitude of TRPV4-mediated currents (at V_m_ = -85 mV) were compared using Students *t*-test. **, P < 0.01; NS: not significant. *N* = 9.

### TRPV4 is predominantly outside of caveolae/lipid rafts

To determine whether TRPV4 is enriched in lipid rafts, detergent-free lipids were isolated by gradient (5%/35%/45% sucrose) ultracentrifugation, and fractions analyzed by Western blot. The 48 kDa lipid raft marker flotillin labeled Fractions 3 - 6. A weak TRPV4 signal was detected in Fraction 4, with the majority of the protein partitioned into the flotillin-free Fraction 2 (Fig. 9A).

**Figure 9.**
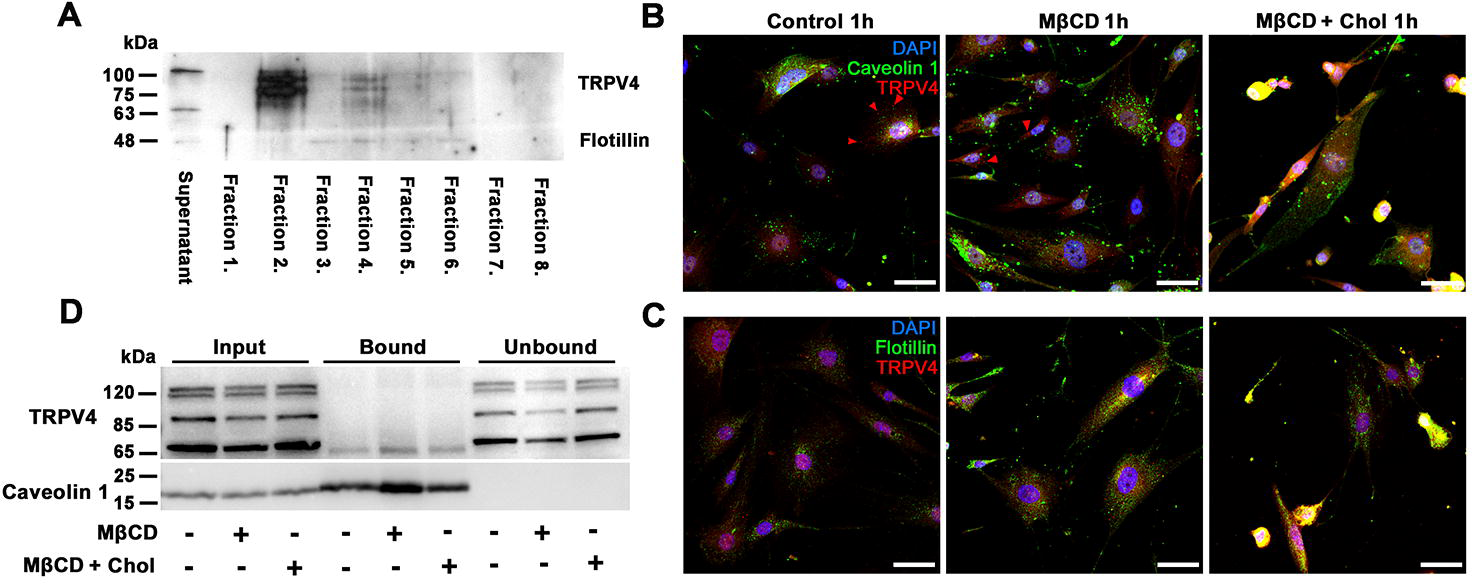
The majority of membrane TRPV4 does not partition into raft domains or interact with caveolar proteins. (A) Western blot; detergent-free lipid raft isolation in primary TM cells. The supernatant fraction contains cytosolic protein; Fractions 1 and 2 (5% sucrose) contain non-lipid-raft membrane protein, Fractions 3-6 (5 - 35% sucrose) contain lipid raft membranes, Fractions 7 and 8 (pellet, 45% sucrose) contain unsuspended protein and cell nuclei. TRPV4 protein is predominantly confined to Fraction 2; a Fraction 4 component associates with flotillin. (B) Co-immunoprecipitation. TRPV4-Cav-1 interaction assessed with Cav-1 antibody for TRPV4 pulldown in control, MßCD and MβCD:cholesterol-treated samples. Input bands, whole cell lysate; Bound bands, Cav-1-bound protein fraction; Unbound bands, flow-through fractions. Cav-1-bound fractions show modest precipitation of the ∼75 kDa TRPV4 isoform. Absence of Cav-1 expression in unbound fractions confirms the quantitative immunoprecipitation of Cav-1. (C & D) Immunohistochemistry, for control, MßCD and CD:cholesterol-treated cells. (C) Double-immunolabeling for TRPV4 and Cav-1. TRPV4-ir β (red arrowheads) does not colocalize with Cav1-ir puncta (green). (D) Double-immunolabeling for TRPV4 and flotillin. TRPV4-ir (red puncta) does not colocalize with flotillin-ir puncta (green). The inset is shown at higher magnification insets as Supplementary Fig. 3.

TRPV4 in cultured endothelial cells coimmunoprecipitates with caveolin-1 (Cav-1; 29; 30). To test the interaction in TM cells, Cav-1 was immunoprecipitated from TM cell lysates and the association of TRPV4 was assessed by Western blotting. The 4 bands in the Western blot (Fig. 9D) presumably correspond to glycosylated, full-length and truncated protein (TRPV4A-E, 56; 57; 58). Following quantitative precipitation of Cav-1, a portion of the ∼ 75 kDa truncated variant associated with Cav-1 whereas all of the full-length variants observed in the total lysate partitioned to Cav-1-depleted, unbound fractions (Fig. 9D). Thus, by far the major proportion of TRPV4 in TM cells localizes to non-caveolar non-raft membrane domains.

Membrane raft/caveolar localization of TRPV4 was further tested with double-labeling immunochemistry. TRPV4-ir puncta (red arrowheads) do not localize into raft domains, as indicated by the absence of colocalization with Cav-1 (Fig. 9B) and flotillin (Fig. 9C). Cholesterol depletion increased the number of Cav-ir puncta without promoting caveolar TRPV4 translocation.

Membrane raft/caveolar localization of TRPV4 was further tested with double-labeling immunochemistry. TRPV4-ir puncta (red arrowheads) do not localize into raft domains, as indicated by the absence of colocalization with Cav-1 (Fig. 9C) and flotillin (Fig. 9D). Cholesterol depletion increased the number of Cav-ir puncta without promoting caveolar TRPV4 translocation.

## Discussion

Cells sense external inputs through activation of specialized transducer molecules embedded within the plasma membrane. In this study we demonstrate that the mechanical milieu regulates the lipid composition of the membrane and that TRPV4, a nonselective cation channel that mediates a wide range of physical and chemical inputs, is highly sensitive to free membrane cholesterol levels. The key findings are (i) TM membrane C/P ratio is regulated by physiological levels of mechanical stretch, (ii) The majority of TRPV4 protein is excluded from raft/caveolar regions, with the possible exception of a small proportion of a truncated splice variant that interacts with Cav-1; (iii) Membrane cholesterol negatively modulates TRPV4 activation by agonists and mechanical stimuli; (iv) TM F-actin expression is cholesterol- and stretch-dependent; and (v) Cholesterol tonically suppresses TRPV4 activity in a non-mammalian expression system. Taken together, these findings implicate cholesterol-TRPV4 interactions in dynamic, use-dependent regulation of cellular mechanosensing.

In contrast to the extensive body of knowledge about how lipid composition affects the structural and mechanical properties of biological membranes, much less is known about the relationship between mechanical stress and the membrane lipid content. Our finding that healthy TM cells respond to physiological levels of cyclic strain with a reduction in the C/P ratio and reduced expression of lipid rafts suggests that the biomechanical environment regulates biophysical and modulatory properties of the membrane, which in turn may fine-tune the function of embedded proteins. Stretch-induced C/P remodeling was inhibited by HC067047, indicating involvement of a TRPV4-dependent step and Δ[Ca^2+^]_i_. Glaucomatous TM cells exhibit elevated C/P ratios (22), suggesting that chronic exposure to elevated IOP might impair the TM capacity to homeostatically compensate for mechanical stress exposure (59; 60; 61).

Free cholesterol is a central constitutent of caveolae and lipid rafts, which serve as portals to regulate cellular cholesterol homeostasis (1; 2; 5; 7; 62). We found that its depletion resulted in near-total dissolution of lipid rafts (visualized by filipin fluorescence), together with increased membrane expression of TRPV4 and Cav-1. MβCD potentiated the amplitude of Ca^2+^ signals induced by the agonist GSK1016790A, cyclic strain and cell swelling and activated transmembrane currents in TRPV4-expressing, but not control, *X. laevis* oocytes, suggesting that cholesterol functions as a negative regulator of TRPV4 and stress fiber signaling in mammalian and nonmmalian systems. Supplementation with exogenous MβCD:cholesterol obviated the facilitation induced by the cyclodextrin alone without affecting peak amplitudes of agonist- and mechanically-evoked Ca^2+^ signals mediated via TRPV4. Activation of the mechanosensitive channel Piezo1 can also be modulated by Mβ CD:cholesterol (69), possibly because control membranes are saturated with free cholesterol, which constitutes ∼third of total membrane lipid mass (5, 35). The ∼4.5-fold increase in the number and intensity of TRPV4-ir puncta (Figure 7) implicates cholesterol in regulation of transport and/or retrieval of TRPV4-containing vesicle pools. MβCD and mutations in cholesterol-binding residues regulate the trafficking of TRPV1 (52, 87), Kv1.5 (85), Kir (86), LRRC8/SWELL (65) and TRPC3 (64) channels. Cholesterol may regulate endocytosis/secretion via PACSIN which binds the N-terminus of the channel (85), SNARE proteins (86) and/or reduced formation of clathrin-coated vesicles (63).

Lipids regulate gating of ion channels through direct interactions within membrane microdomains and indirectly by shaping the biophysical properties of the membrane. Stretch-activated channels might be activated by increased tension caused by cholesterol removal (66, 83), however, viscoelastic “force-from-lipid” models (64, 75, 76) cannot explain why ion channels respond differently to cholesterol enantiomers with similar effects on membrane properties (77). Given that cholesterol depletion facilitates TRPV4 activation in TM cells and oocytes but *suppresses* TRPV4 activation in glia (70) and endothelial cells (29), the most likely model is allosteric modulation of channel subunits, accessory proteins and/or residues buried within the bilayer. Consistent with this, MβCD facilitates mammalian TRPM8 (50), TRPM3 (51), TRPC3 (64) and VRAC (volume-activated chloride) channel activation (65) while inhibiting TRPA1, TRPC1, TRPC6 channels (71–73), and blocking TRPL currents in fly photoreceptors (74). Allosteric sites in vanilloid TRP isoforms may include Cholesterol Recognition/Interaction Amino acid Consensus (CRAC)-like (KDLFRFLL) recognition motifs that span Loop 4 – TM5 (28, 82, 84). Consistent with our results, loss of TRPV4-cholesterol interaction in the TRPV4^R616Q^ CARC mutation was associated with gain-of-function for TRPV4 (67). It remains to be seen whether cholesterol influences the context of the cellular sensory response. For example, TRPV4 activity in yeast (which cannot synthesize cholesterol) can be induced by swelling but not temperature (79) whereas the mammalian channel shows a pronounced activation optimum at ∼34-28 °C. In addition, MβCD inhibited capsaicin- and proton-evoked currents (80, 81) but facilitated thermal sensitivity of the cognate TRPV1 channel (80).

Mechanosensing often involves interactions between the cell membrane and the cytoskeleton. TRPV4 regulates cytoskeletal dynamics through direct interactions and Ca^2+^- dependent signaling (68), with TM cells compensating for membrane stretch by increasing stiffness through TRPV4-dependent increase in the density of stress fibers and focal complexes CD treatment dissolves lipid rafts and stimulates actin polymerization is in accord with the reports in osteoblasts, fibroblasts, endothelial cells, and myocytes (88–90) and links the effects of MβCD to downstream increases in RhoA activation, contractility and increased cell stiffness (90). The additivity of stretch and cholesterol depletion (Figure 6C) suggests that stretch-activated channels and cholesterol-regulated membrane domains control independent signaling mechanisms that remain to be elucidated in future work.

In contrast to direct cholesterol depletion by MβCD, simvastatin, which blocks synthesis of the cholesterol precursor mevalonate, did not affect the amplitude of stretch-induced Ca^2+^ signals.

Differential effects of simvastatin vs. MβCD on stress fiber formation and cell contractility were also observed in endothelial cells and fibroblasts (78; 88).

Outflow of aqueous humor is fine-tuned by arrays of mechanosensitive molecules that include lipid rafts, caveolae, focal complexes, as well as TRPV4, Piezo1 and TREK-1 channels (24; 61; 91; 92; 93; 98). Many TRP isoforms were reported to traffic into lipid rafts (72), including TRPV4 which was suggested to interact with Cav-1 in endothelial and smooth muscle cells (29; 67; 30). Because both TRPV4 and Cav-1 contribute to TM mechanosensitivity and outflow homeostasis (24; 25; 94; 95; 96), we expected TRPV4 expression to be confined within cholesterol-enriched raft/caveolar domains. This turned out not to be the case. TRPV4 did not colocalize with filipin (lipid raft marker), Cav-1 (marker of caveolae) and flotillin (marker of non-caveolar rafts), and the majority of the TRPV4 protein partitioned into fractions that excluded raft proteins. The human TRPV4 protein consists of 871 amino acids with at least five variants with unknown differences in function (97; 58). The four bands seen in TM, kidney, ventricular myocyte and choroid epithelial immunoblots (56; 57; 58; Figure 9) correspond to glycosylated full-length variants and an unglycosylated ∼75 kDa splice variant of TRPV4. By far the greatest fraction of the protein, including all high-M.W. variants, partitioned into total lysate/unbound fractions that did not contain Cav-1/flotillin and resisted cholesterol depletion (Fig. 9D). Interestingly, Cav-1 precipitated a small portion of the ∼75kDa variant, suggesting that TRPV4 variants might be differentially susceptible to caveolar interactions. Non-raft low-cholesterol regions reduce membrane mobility of TRPV4, with loss of cholesterol interaction resulting in gain-of-function for the channel (67). Similar results were observed for TRPM8, which shows enhanced gating in non-raft domains (50). Future work will show whether TRPV4 variants differ in microdomain interactions with lipids and the role of specific TRPV4 variants in allosteric modulation by raft-enriched G-proteins, protein kinases/phosphatases and Ca^2+^-binding proteins.

In summary, our study suggests that TM cholesterol homeostasis is integrated with mechanotransduction to regulate multiple processes relevant for IOP regulation. Cholesterol may affect cellular function both by altering bulk properties of the membrane (steady-state tension, lipid composition, stiffness) and through allosteric modulation of resident proteins. The interdependence between membrane tension, membrane cholesterol content, TRPV4 signaling and calcium homeostasis points at a dynamic mechanism that may protect TM cells and the conventional outflow pathway from hypertension-induced injury yet lose adaptive function in diseases involving chronic increases in mechanical stress or pathological dysregulation of cholesterol metabolism such as glaucoma, diabetic retinopathy, Niemann-Pick disease and/or macular degeneration.

## 2. MATERIALS AND METHODS

### TM cell culture and isolation

Primary cultures of TM (pTM) cells were dissected from 3 eyes of donors with no history of eye disease (65-year-old male, 68-year-old male, 78-year-old male) as described previously (24, 25, 98) and in concordance with the tenets of the WMA Declaration of Helsinki and the Department of Health and Human Services Belmont Report. A subset of experiments was conducted in immortalized juxtacanalicular human trabecular meshwork (hTM cells) obtained from ScienCell (Catalog#6590) and used up to the 7^th^ passage. pTM and hTM cells showed no differences in responses to TRPV4 or cholesterol-modulating agents. Cells were grown in Trabecular Meshwork Cell Medium (TMCM, ScienCell, Catalog#6591) at 37 °C and L 5% CO_2_. The phenotype was periodically validated by profiling for markers, including *Aqp1*,α SMA) and dexamethasone-induced upregulation of myocilin expression. These data are shown in our previous characterizations of the cell line (21, 24, 98).

*Xenopus laevis* oocyte experiments were performed according to the guidelines of the Danish Veterinary and Food Administration (Ministry of Environment and Food), and approved by the animal facility at the Faculty of Health and Medical Sciences, University of Copenhagen. The experiments conform to the principles and regulations described in (99). The surgical protocol by which the oocytes were retrieved was approved by The Danish National Committee for Animal Studies, Danish Veterinary and Food Administration (Ministry of Environment and Food). An abstract version of this work appeared in (100).

### Reagents

The TRPV4 agonist GSK1016790A (GSK101) and cholesterol were obtained from Sigma or VWR. GSK101 (1 mM) stock aliquots were prepared in DMSO and subsequently diluted into working saline concentrations (5 and 25 nM, respectively). Chemical reagents for biochemical experiments - methanol, isopropanol, n-hexane, and chloroform - were of GC/MS grade and purchased from Fisher Scientific. The cholesterol standard was purchased from Sigma-Aldrich.

### Cholesterol depletion and repletion

Methyl-β-cyclodextrin (MβCD; Sigma C4555) was dissolved in TMCM and used at 10 mM, the concentration that removes 80-90% of free membrane cholesterol (32, 35). Cells were preincubated with MβCD for 60 min to maximize extraction, washed in TMCM and placed into recording chambers for optical recordings. This protocol maintains decreased membrane cholesterol levels for at least 24 hours (86).

Supplementation was based on perfusion with cholesterol-saturated MβCD. Powdered cholesterol was dissolved for 30 min in 80% ethanol solution at 75 - 80° C to obtain a 10 mM stock solution. The stock was dissolved in TMCM containing M CD to obtain the final cholesterol concentration of 1 mM. A parallel chamber used the cholesterol stock to load MβCD for the final concentrations of 1 mM cholesterol + 10 mM MβCD.

### Hypotonic stimulation

The swelling studies were conducted as reported previously (42, 46). Extracellular NaCl was kept at 57.5 mM and total osmolarity regulated by addition or removal of mannitol, a procedure that maintains the ionic strength of the extracellular solution. Osmolarity was checked thermometrically using a vapor pressure osmometer (Wescor).

### Lipid extraction and GC-MS Chromatography

Lipids were extracted using Folch method (101). Total lipids were extracted by adding methanol/chloroform/water (1:2:1, v/v). The chloroform phase was washed to remove water residues and dried under nitrogen gas. The dried film was dissolved in 100 µl hexane and 5 µl of sample was injected into the GC-MS instrument for cholesterol analysis. The Thermo Trace GC-DSQ II system (ThermoFisher Scientific) consists of an automatic sample injector (AS 3000), gas chromatograph (G.C.), single quadrupole mass detector, and an analytical workstation. Chromatographic separation was carried out with an Rxi- 5MS coated 5% diphenyl/95% dimethyl polysiloxane capillary column (30 m × 0.25 mm i.d, µm film thickness) (Restek Corporation, PA). The sample was injected into the GC/MS using a splitless mode, the septum purge was on, and the injector temperature was set at 200°C. The column temperature was programmed as follows: initial temperature 60°C, 15 degrees/min to 240°C, 2 degrees/min to 290°C, and a hold at 290°C for 5 min. Transfer line temperature was 290°C. Helium was used as the carrier gas at a flow rate of 1.5 ml/min. M.S. conditions were as follows: electron ionization mode with ion source temperature of 250°C and multiplier voltage, 1,182 V; full scan and selected ion monitoring (SIM) mode was used for the identification and quantification of cholesterol. The area values of cholesterol were plotted against known range of standards (100 to 0.1 ng) to quantify the cholesterol in the samples. Phosphatidylcholine levels were measured by a colorimetric/fluorometric assay kit (BioVision).

### Immunprecipitation and Western blot analysis

Cell lysis was performed in lysis buffer containing 2% octylglucoside, 150 mM NaCl, 10 mM Tris-HCl pH 7.4, 0.5 mM EDTA, 0.1% Triton X-100 and protease inhibitor cocktail (Roche). Lysates were cleared by centrifugation and protein concentrations determined using a BCA assay (ThermoFisher Scientific). Immunoprecipitation of Cav-1 was performed using Dynabeads™ Protein G Kit (ThermoFisher Scientific) according to the manufacturer’s instructions. Briefly, Cav-1 primary antibody (10 ng/µl) was conjugated to Dynabeads magnetic beads at room temperature (R.T.). Then, equal amount of proteins from each sample were incubated with the beads-antibody conjugates for 15 min at R.T., with gentle agitation. The beads were removed from solution by DynaMag™-2 magnet (ThermoFisher Scientific), washed in wash buffer and resuspended in lysis buffer. The original cell lysates, immunoprecipitates and unbound fractions (flow through) were boiled in Laemmli buffer, separated by reducing SDS-PAGE and transferred to nitrocellulose membranes. Membranes were blocked with 5% bovine serum albumin (BSA) for 1 h, and probed with Cav-1 (Cell Signaling Technology) and TRPV4 primary antibodies (Alomone Lab). Primary antibodies were detected using Clean-Blot^®^ IP detection reagent (ThermoFisher Scientific) conjugated to horseradish peroxidase (HRP).

### Detergent-free lipid raft isolation and Western blot analysis

Samples were homogenized in hypotonic homogenization buffer (20mM Tris-HCL pH 7.8, 3 mM MgCl_2_ 10 mM NaCl, 0.0005 mg/ml, 2 mM Na-vanadate, 20 mM Na-fluoride, 0.5 mM DTT, 1mM PMSF) on ice and centrifuged at 15,000 g for 30 min at 4 C° to separate cytosolic proteins from intracellular and plasma membranes. The pellet was resuspended in 0.5 M Na_2_CO_3_, transfered to a 5%/35%/45% sucrose (in Na_2_CO_3_) flotation-gradient and spun at 36,000 rpm for 18 hrs using a preparative ultracentrifuge model XL-90 (NVT90 rotor; Beckman Coulter Life Sciences). Fractions obtained from the sucrose gradient were diluted in hypotonic buffer and spun ar 15,000 g for 30 min at 4 C°. Pellets (25 µl) were resuspended in RIPA buffer and 2x Laemmli buffer. 30 µl of each sample was loaded in 10% SDS-PAGE, and transferred to polyvinylidene difluoride (PVDF) membranes for 1h at 220 mA. Non-specific binding was blocked with 5% non-fat milk and 2% BSA. The samples were incubated overnight at 4 °C with TRPV4 (1:500, Alomone Labs) and flotillin (1:200, Santa Cruz Biotechnology) antibodies, followed by anti-mouse (1:5000, BioRad) or anti-rabbit (1:5000, Cell Signaling) horseradish peroxidase (HRP)-conjugated secondary antibodies. The blotted proteins were developed with an enhanced chemiluminescence kit (Thermo Fisher Scientific).

### Immunofluorescence

Cells were fixed with 4% paraformaldehyde for 10 min. After a PBS rinse, PBS containing 5% FBS and 0.3% Triton X-100 blocking solution was applied for 20 minutes. F-actin was labeled with AlexaFluor 488 phalloidin (1:1000, Life Technologies). Primary antibodies (rabbit anti-TRPV4, 1:1000, Lifespan Biosciences, mouse anti-flotillin, 1:200, Santa Cruz; mouse anti-caveolin, 1:1000, B.D. Biosciences) were diluted in antibody solution (2 % BSA, 0.2 % TritonX-100 in PBS) and applied overnight at 4°C. The TRPV4 antibody does not label TRPV4 KO tissues (102, 103). After rinsing, slices were incubated with secondary antibodies diluted to 1:1000 in PBS for one hour at R.T. Plasma membrane cholesterol was tracked with filipin (Sigma F9765). As previously described (70), 0.005% filipin (Sigma) was dissolved in DMSO and applied to dissociated cells together with the secondary antibody (goat anti-rabbit AlexaFluor 488; 1:500; Life Technologies). Unbound antibodies were rinsed and conjugated fluorophores protected with Fluoromount-G (Southern Biotech) prior to coverslipping. Images (10 per experiment) were acquired on Olympus CV1200 confocal microscope using a NeoFluor 20x water immersion objective.

### Analysis and particle counting

Images were acquired using identical parameters (H.V., gain, offset) resulting in very similar signal-to-noise ratios across datasets. ImageJ was used to extract and quantify the mean intensities and particle analysis of immunoreactive signals, with ∼40-50 cells per slide averaged across at least 3 independent experiments. The fluorescence intensity of F-actin was measured in arbitrary units using the Area Integrated Intensity measurement tool of ImageJ with background compensation. Data are plotted the signal as averaged and normalized florescence intensity (in %) per cell area compared with the control. In particle analysis, color images were converted to black and white using *Binary*

*Convert to mask* with white background and automatic threshold level. Immunoreactive puncta (number/cell area) with the segmented area were counted with the *Analyze particles* plug-in. Minimum (3 pixel^2^) and maximum (30 pixel^2^) pixel area sizes were defined to exclude areas outside of the ROI, calculate the particle number/cell area and determine the relative particulum numbers. Individual particulum sizes for each cell were averaged and normalized.

### Optical Imaging

Calcium responses in TM cells were tracked following published protocols (102, 104). Briefly, for CCD imaging, cells were loaded with 5-10 µM Fura-2-AM for 45 min and perfused with isotonic saline (pH 7.4) containing (in mM): NaCl 133, KCl 2.5, NaH_2_PO_4_ 1.5, MgCl_2_ (6H_2_0) 1.5, CaCl_2_ 2, glucose 10, HEPES hemisodium salt 10, pyruvic acid 1, lactic acid 1, L-glutamine 0.5, glutathione 0.5, Na-ascorbate 0.3, with pH 7.4 and osmolarity at 300 ± 10 mOsm, delivered through a gravity-fed 8-reservoir system (Warner Instruments) that converged towards a manifold tube inserted into the experimental chamber. Epifluorescence was detected with Photometrics Delta or Prime BSI cameras. Ratiometric Ca^2+^ imaging was performed on Regions of Interest (ROI) that marked a central somatic region, and were typically binned at 3x3 (102, 104). Background fluorescence was measured in similarly sized TM ROIs in neighboring areas devoid of cells. The microscopes were inverted Nikon Ti with 40x (0.75 NA oil) or upright Nikon E600 FN microscopes with a 20x (0.8 NA water) and 40x (1.3 N.A. oil & 0.8 N.A. water) objectives. A wide-spectrum 150W Xenon arc lamp (DG4, Sutter Instruments, Novato, CA) provided excitation to 340 nm and 380 nm filters (Semrock). The signals were analyzed using NIS-Elements AR 3.2 and M.S. Excel ΔR/R (peak F^340^/F^380^ ratio –baseline/baseline) was used to quantify the amplitude of Ca^2+^ signals. Data acquisition and F340/F380 ratio calculations were performed by NIS Elements 3.22 software (Melville, NY).

### Spectrophotometry

[Ca^2+^]_i_ was monitored using a plate reader (Turner Biosystems). Cells were seeded onto non-coated 96-well plates and loaded with 2 uM of Fluo-4 AM for 30-45 min at 37 C. Fluorescence was measured at 490 nm excitation and 520 nm emission, with 90 sec L intervals, and 6-10 measurements per experiment. 490/520 nm ratios were normalized to the control untreated samples. Baseline measurements were recorded in control wells at the same time withoutaddition of agonist or HTS.

### Membrane strain assay

TM cells were seeded on flexible silicon membranes coated with type I/IV collagen, grown to 80% confluence and placed into a FlexJunior chamber controlled by the Flexcell-5000 Tension system (Flexcell) (24). For determination of stretch-induced Ca^2+^ influx and cytoskeletal changes, cells were loaded with Fura-2-AM for 30-60 minutes and stimulated with cyclic biaxial stretch (10%, 1Hz, 15 min, or 6%, 0.5Hz 1-3 hours; respectively) at 37 C. A L 15 minute stretch period was chosen to optimize capture of the stretch-evoked fluorescent response by adjusting for the change in focal plane (which disrupted calcium imaging for several seconds as indicated by breaks in response trace in Fig. 5A). Cells were imaged with a Nikon E600FN upright microscope. Excitation light was provided by a Xenon source within a Lambda DG4 (Sutter Instruments) and controlled by Nikon Elements.

### RNA preparation and heterologous expression in Xenopus laevis oocytes

*Xenopus laevis* frogs were obtained from Nasco (Fort Atkinson). The frogs were kept in tanks in a recirculating water facility, and fed twice weekly with Floating Frog Food 3/32 for adults and juveniles (Xenopus Express, Inc.). Oocytes were surgically removed from anesthetized frogs (40, 46) under anesthesia (2 g/L Tricain, 3-aminobenzoic acid ethyl ester, Sigma A-5040). Preparation of defolliculated oocytes was carried out as described in (106). Oocytes were kept in Kulori medium (in mM): 90 NaCl, 1 KCl, 1 CaCl_2_, 1 MgCl_2_, 5 HEPES (pH 7.4). cDNA encoding rat TRPV4 was subcloned into the oocyte expression vector pXOOM, linearized downstream from the poly-A segment, and *in vitro* transcribed using T7 mMessage machine according to manufacturer’s instructions (Ambion). cRNA was extracted with MEGAclear (Ambion) and microinjected into defolliculated *Xenopus laevis* oocytes: 4 ng TRPV4 RNA/oocyte. A TRP channel antagonist, Ruthenium Red (1 μM, Sigma Aldrich, R-2751) was added to the medium to μ prevent the tonic TRPV4-mediated currents mediating oocyte lysis. The oocytes were kept for 3- 4 days at 19°C prior to experiments.

### Electrophysiology on Xenopus laevis oocytes

Conventional two-electrode voltage clamp studies were performed with a DAGAN CA-1B High Performance oocyte clamp (DAGAN) with Digidata 1440A interface controlled by pCLAMP software, version 10.5 (Molecular Devices). Electrodes were pulled (HEKA, PIP5) from borosilicate glass capillaries to a resistance of 2.5-3 MΩ when filled with 1 M KCl. The current traces were obtained in a test solution containing (in mM): 50 NaCl, 2 KCl, 1 MgCl_2_, 1 CaCl_2_, 10 HEPES, 100 mM mannitol (Tris buffered pH 7.4, 220 mOsm) by stepping the clamp potential from -20 mV to test potentials ranging from +50 mV to -130 mV (pulses of 200 ms) in increments of 15 mV. Recordings were low pass-filtered at 500 Hz, sampled at 1 kHz, and the steady-state current activity was analyzed at 140-180 ms after applying the test pulse. Depletion of endogenous cholesterol in intact oocytes was induced with 50 μM MβCD for 45 min (e.g., 55). All experiments were performed at room temperature (23 °C).

### Data Analysis

Statistical analyses were performed with GraphPad Prism 6.0 and Origin Pro 8.5. Data was acquired from at least three different experimental preparations on different days, with 3-6 slides/experiment. Typically three different batches of oocytes were used. Unless indicated otherwise, unpaired or paired t-tests were used to compare two means, and a one-way ANOVA along with the Tukey’s test was used to compare three or more means. Means are shown ± SEM. p > 0.05 = NS, p < 0.05 = *, p < 0.01 = **, p < 0.001 = *** and p < 0.0001 = ****.

### Data Availability

The data described in the mansucript are contained within the manuscript.

## Acknowledgements

We thank Dr. Peter Geck (Tufts University, Boston MA) for the generous help with lipid raft experiments.

---

**Supplementary Figure 1.** Averaged, maximum fluorescence data in Fluo-4 loaded TM cells. (A) Spectrophotometry shows dose-dependent [Ca^2+^]_i_ increases in response to 5 and 25 nM GSK and augmentation by cholesterol removal (N = 4) (B) Data shows dose-dependent [Ca^2+^]_i_ increases in response to different HTS solutions and augmentations by cholesterol removal (N = 3). *, P < 0.05.

**Supplementary Figure 2. Membrane cholesterol modulates cyclic stretch induced [Ca^2+^]_i_ response.** (A) Representative traces and (B) Averaged results for 15 min cyclic stretch of control β CD:cholesterol-treated (green trace, bar) and simvastatin (1mM, red trace, bar)-treated cells. Stretch-evoked [Ca^2+^]i responses are facilitated by Mβ CD:cholesterol (green bar) decreased the amplitude of the response. Simvastatin (1mM) did not facilitate the stretch effect (n = 21-77; N = 5-6); ****, P < 0.0001.

**Supplementary Figure 3 - The majority of membrane TRPV4 does not partition into raft domains or interact with caveolar proteins.** Higher magnification insets of images Fig. 9C & D.

