## Supplementary figures and images for "Membrane cholesterol regulates TRPV4 function, cytoskeletal expression, and the cellular response to tension"

### Supplemental Figure 1

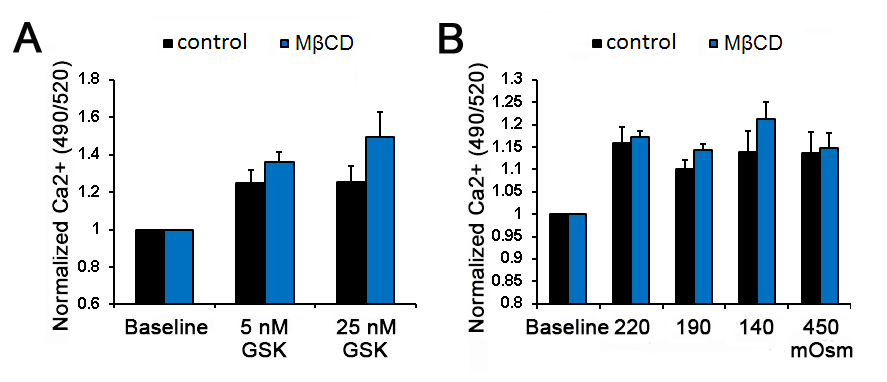

### Supplemental Figure 2

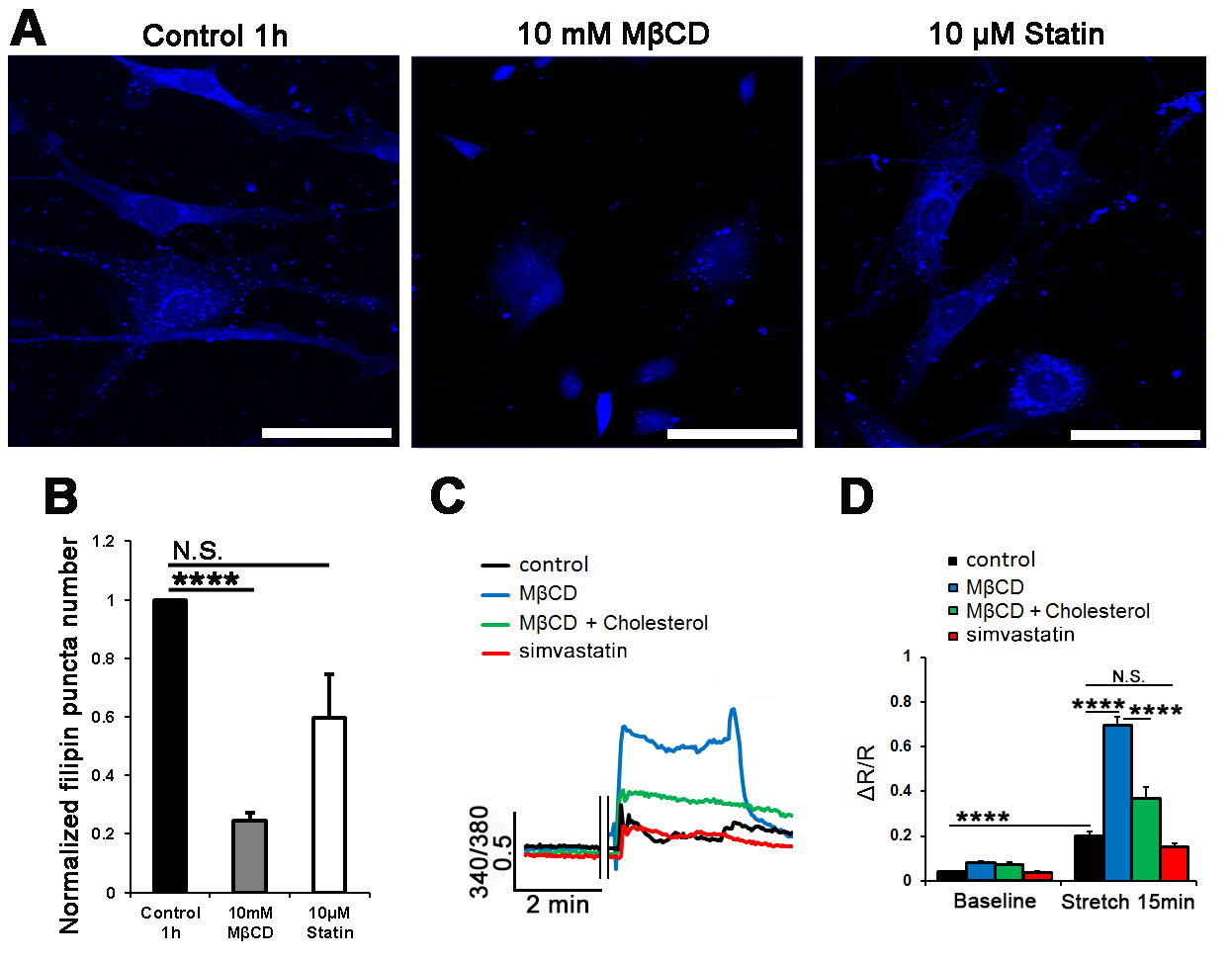

### Supplemental Figure 3

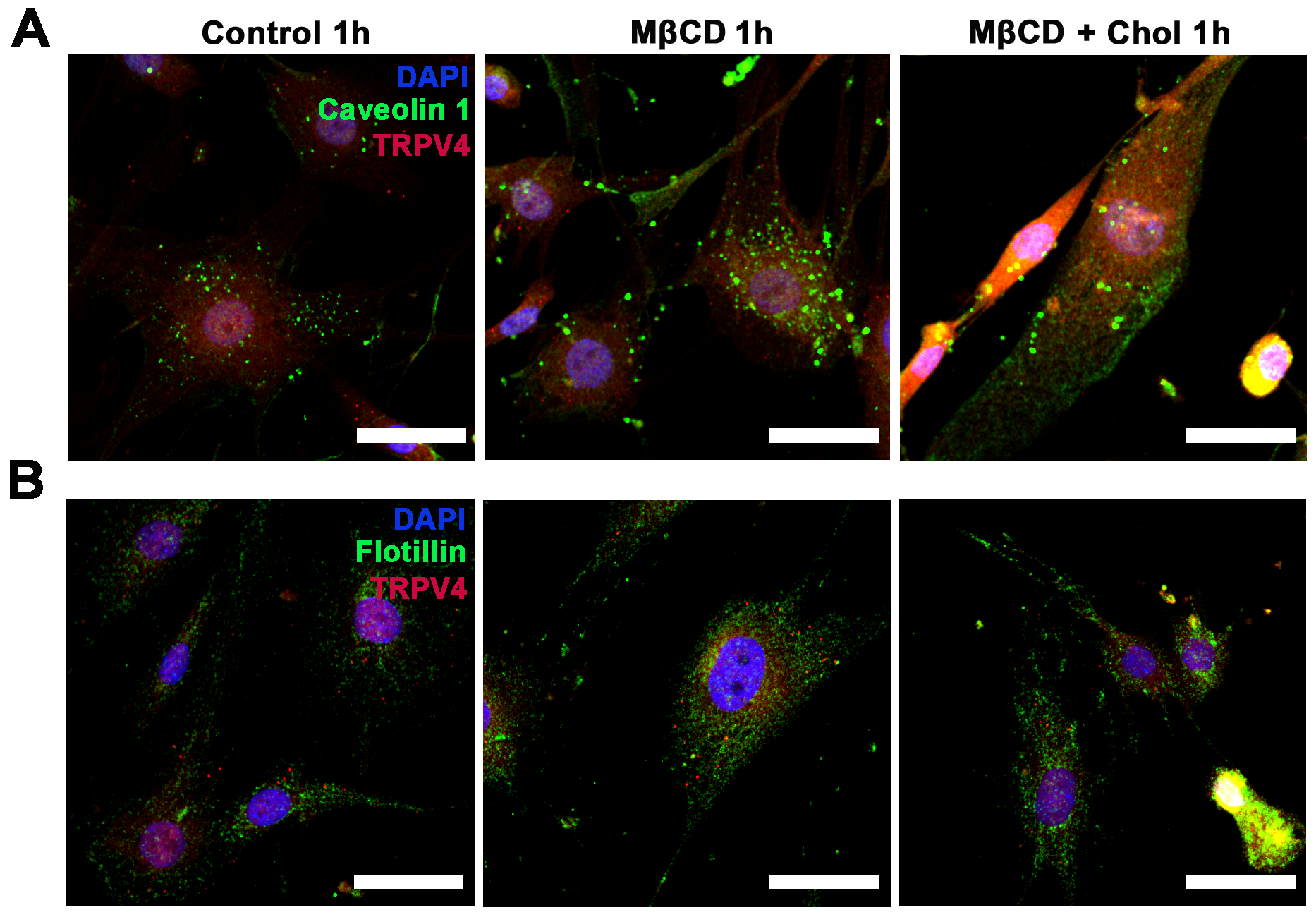
